# Optimizing High-Parameter T-Cell Immunophenotyping Through Direct Comparison of Conventional and Spectral Flow Cytometry

**DOI:** 10.64898/2026.04.17.718631

**Authors:** Domenico Lo Tartaro, Kelly Lundsten, Anisha Jose, Andrea Cossarizza

## Abstract

High-parameter flow cytometry is essential for dissecting the intricate landscape of T-cell diversity. In this study, we directly compare conventional flow cytometry (CFC) and spectral flow cytometry (SFC) for high-dimensional T-cell phenotyping, assessing how spectral detection and panel-design strategies influence analytical performance.

Using peripheral blood mononuclear cells from healthy donors stained with both an established (v1) and an optimized (v2) fluorochrome-labelled antibody panel, and analyzed through manual gating and unsupervised approaches, we found that CFC reliably identified major T-cell subsets. However, spectral acquisition consistently delivered clear technical advantages, including improved signal-to-noise ratios, higher staining index values, and superior resolution of low-intensity and co-expressed markers.

These improvements translated into more sharply delineated multidimensional clusters and a markedly enhanced resolution of T-cell differentiation states. Moreover, the optimized spectral panel enhanced the unsupervised detection of rare populations, such as cytotoxic CD4⁺ T-cells (PD-1⁺GZMB⁺). However, despite the overall increase in data quality achieved with SFC, the selection of antibody clones may influence the measured frequencies of the identified populations. Finally, SFC - particularly when coupled with rational panel optimization and the use of advanced fluorophores - consistently delivers superior, higher-quality measurements and improved multidimensional resolution, thereby substantially enhancing the robustness and sensitivity of high-parameter T-cell phenotyping for comprehensive immunological studies.

## Background

T-cells are central effectors of adaptive immunity, orchestrating protection against pathogens and tumors, whereas dysregulated T-cell responses drive chronic inflammation and autoimmune pathology. Their functional diversity and dynamic differentiation states require analytical methods capable of high-resolution phenotyping. Flow cytometry remains a cornerstone for characterizing T-cell lineages, differentiation stages, activation and exhaustion programs, and tissue-resident populations in both healthy and disease contexts.^1–5^

Recent advances in high-parameter cytometry have expanded conventional panels from 8–12 colors to 30–50 markers, enabling detailed interrogation of memory subsets, T follicular helper (Tfh) and regulatory T-cell (Treg) populations, chemokine-receptor landscapes, and checkpoint regulation, all with minimal sample input.^6–8^ Spectral flow cytometry further extends this capacity by resolving full emission spectra, allowing for improved separation of co-expressed markers, expanded fluorochrome selection, and deeper phenotyping of rare or functionally distinct populations.^9^ Structured spectral panel design emphasizes optimized single-stain references, rigorous unmixing validation, autofluorescence correction, and longitudinal stability, facilitating reproducible high-dimensional assays suitable for immune monitoring and multi-site studies.

Conventional flow cytometry continues to provide a robust foundation for validated T-cell panels; particularly when panel sizes are moderate, compensation-based workflows are preferred, or harmonization with legacy datasets is required.^10^ Established panel structures, fluorochrome combinations, and gating strategies ensure consistency, reproducibility, and regulatory familiarity across translational and clinical studies.

Translating high-parameter panels from conventional to spectral platforms raises several technical considerations. Marker resolution and population frequencies must be evaluated across platforms, fluorochrome reassignment may be required to reduce spreading error and optimize co-expression resolution, and spectral workflows require robust unmixing controls, especially for dim markers or tandem dyes.^4^

We evaluated a comprehensive T-cell panel including markers for lineage (CD3, CD4, CD8), differentiation (CCR7, CD45RA), tissue residency (CD69, CD103, CD49a), checkpoint receptors (PD-1, CTLA-4, CD39, CD96), co-stimulatory receptors (CD226), chemokine-guided trafficking (CXCR6), and functional/proliferative status (T-bet, Granzyme B [GZMB], Ki-67).^11^ The panel was implemented in two formats: an established conventional version (v1) and an optimized spectral version (v2) in which fluorochromes were reassigned to improve resolution and minimize spreading error while retaining biological targets and largely identical clones.

Here, we directly compare conventional and spectral flow cytometry for high-parameter T-cell panels, integrating both manual and unsupervised analyses. Our goal is to evaluate reproducibility, fidelity, and overall analytical performance across platforms while providing practical guidance for translating conventional panels into spectral workflows with high-dimensional interpretability.

## Results and discussion

Peripheral blood mononuclear cells (PBMCs) from five healthy donors (30-65 years old) were used throughout the study. Cells were stained with the original antibody panel (v1),^11^ acquired in both conventional detection mode (conventional flow cytometry; CFC) or in spectral mode (spectral flow cytometry; SFC), or with an optimized spectral panel (v2), acquired exclusively in spectral mode (**Fig. 1A**). Panel optimization involved selective clone substitution and fluorochrome reassignment to improve spectral compatibility and marker resolution (**Supplementary Fig. 1**).

**Figure 1.**
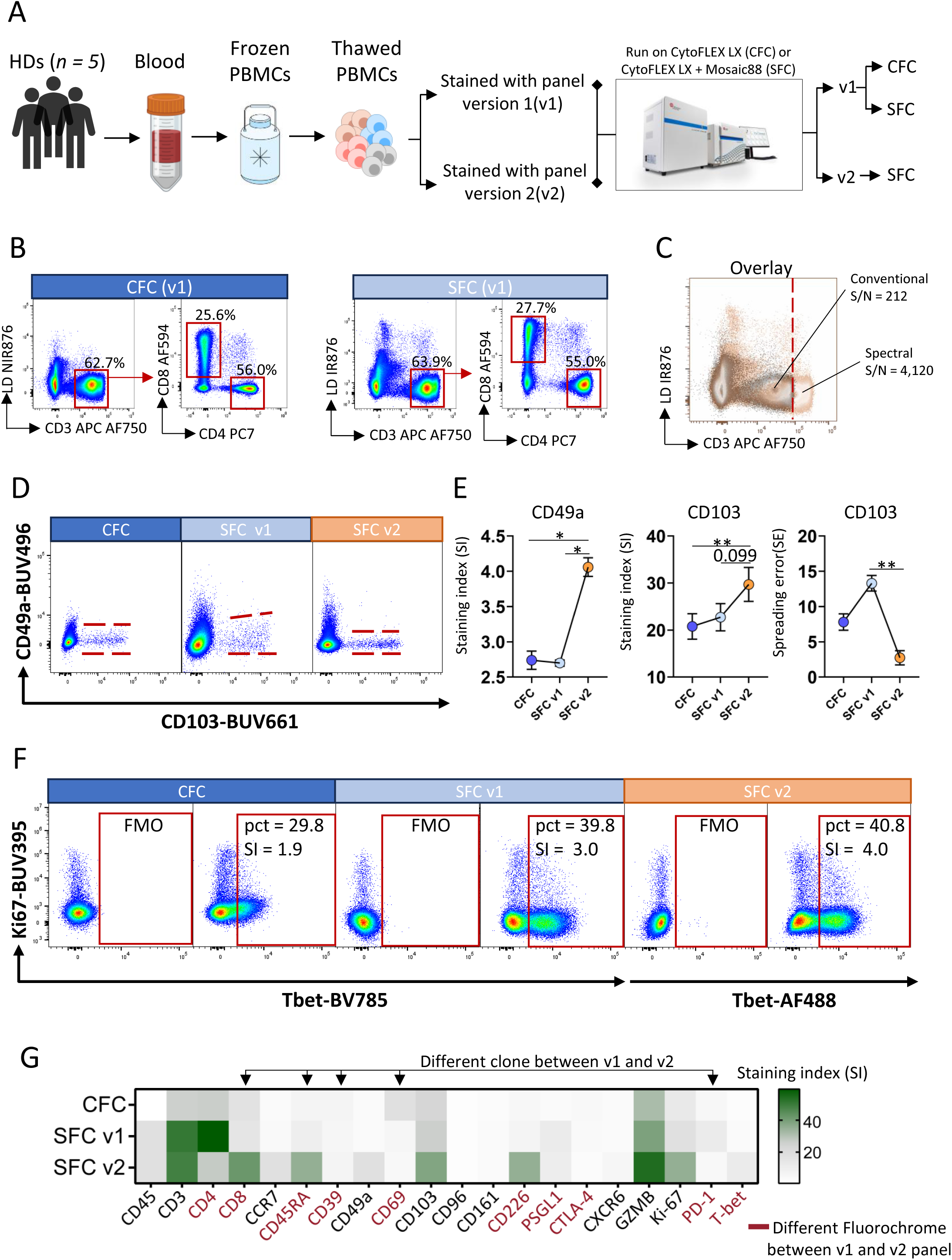
Manual analysis of cryopreserved PBMCs acquired by conventional (CFC) and spectral flow cytometry (SFC). (**A**) Schematic representation of the experimental workflow. Analyses were performed on peripheral blood mononuclear cells (PBMCs) obtained from a healthy donor. PBMCs were stained with either the original antibody panel (v1) or the optimized spectral panel (v2) and acquired using a CytoFLEX LX for conventional flow cytometry (CFC) or a CytoFLEX LX flow cytometer equipped with the CytoFLEX mosaic 88 Spectral Detection Module for spectral flow cytometry (SFC). (**B**) Dot plots of live CD3⁺ T-cells and CD4⁺ and CD8⁺ T-cell subsets stained with the same antibody panel (v1) and acquired using either conventional or spectral detection modes. (**C**) Overlay of CD3⁺ live cells acquired by CFC or SFC v1. The thick red line indicates the upper boundary of CD3 signal in the CFC dataset. Signal-to-noise (S/N; calculated as MFI^+^÷MFI^−^) values for CD3 in both CFC and SFC are reported. (**D**) Spillover spreading error (SE) between CD49d-BUV496 and CD103-BUV661 in CD4⁺ T-cells in CFC, SFCv1, and SFCv2. The red lines indicate the threshold of positivity in the BUV496 channel according to BUV661 positivity. (**E**) Staining index (SI) and standard error (SE) values for CD49a and CD103 in CFC, SFCv1, and SFCv2 are presented as mean ± SEM. A detailed description of SI and SE calculations is provided in the Methods section. Statistical analyses were conducted using the Friedman test, followed by Benjamini–Hochberg correction to control the false discovery rate (FDR). Statistical significance was defined as FDR < 0.05 (*) and FDR < 0.01 (**). (**F**) Representative dot plots showing T-bet and Ki-67 expression in CD8⁺ T-cells acquired by CFC, SFCv1, and SFCv2. Fluorescence-minus-one (FMO) controls were used to define positivity; numbers indicate the percentage and SI of gated positive cells. (**G**) Heatmap summarizing SI values for all markers (except Live/Dead NIR876) across CFC, SFCv1, and SFCv2. Antibodies for which the fluorochrome was changed between v1 and v2 are shown in red; those highlighted in red with a black arrow on the top indicate changes in both fluorochrome and clone between panel versions.

To compare panel performance across detection strategies, major T-cell lineage populations^5,10^ were first evaluated using PBMCs stained with panel v1 and acquired by CFC or SFC. Manual gating revealed that the frequencies of total CD3⁺, CD4⁺, and CD8⁺ T-cells were generally consistent between platforms, indicating comparable identification of primary populations (**Fig. 1B** and **Supplementary Fig. 2**). However, subtle differences were observed in the proportion of total CD8⁺ T-cells, primarily due to variations in the antibody clone and fluorophore employed, as well as low CCR7 signal intensity, which contributed to underestimation of Tn and Tcm subsets (**Supplementary Fig. 2A**). Notably, spectral acquisition greatly enhanced the CD3 signal-to-noise ratio (approximately 212 by CFC *vs.* 4,120 by SFC), demonstrating the increased sensitivity and improved signal resolution afforded by full-spectrum detection (**Fig. 1C**).

Fluorochromes with partially overlapping emission spectra introduce spillover spreading error after compensation or unmixing, which can impair resolution when markers are co-expressed.^7^ Using SFC, we found that this approach improved CD103-BUV661 discrimination and staining index relative to CFC, but increased spread in v1, potentially limiting accurate identification of double-positive populations in complex samples (**Fig. 1D–E**). The optimized spectral panel (v2), incorporating targeted fluorochrome reassignment, further enhanced marker resolution and reduced spreading, highlighting the value of fluorochrome optimization in spectral workflows (**Fig. 1D–E**). These findings are consistent with prior reports showing improved separation of overlapping fluorochromes via panel optimization and full-spectrum unmixing.^9^

Given the challenges of intracellular staining,^12^ including signal attenuation, increased background, and fixation/permeabilization variability, we examined T-bet-BV785 expression within CD8⁺ T-cells as a representative intracellular marker^13^ (**Fig. 1F**). Spectral acquisition improved positive–negative separation and increased the staining index (SI) from 1.9 (CFC) to 3.0 (SFC v1), with a corresponding increase in T-bet⁺ cells (29.8% to 39.8%). Reassigning T-bet to AF488 in SFC v2 further increased the SI to 4.0 without affecting population frequency (40.8%). These results indicate that optimized fluorochrome selection enhances resolution of low-intensity intracellular markers while preserving biological interpretation.

A global comparison of staining indices confirmed progressive improvements from CFC to SFC v1 and additional gains in SFC v2 (**Fig. 1G**), particularly for markers with reassigned fluorochromes or clones (e.g., CD8, T-bet, CD226). While overall cell frequencies remained stable, SFC detection strategies (but not fluorochrome reassignment) contributed to the observed increase in T-bet levels compared with CFC (**Supplementary Fig. 3A**). Notably, PD-1 displayed higher SI and a greater frequency of positive cells in SFC v1. This difference was largely attributable to antibody clone variation rather than fluorochrome brightness when compared with SFC v2 (Δ% = −7.3), highlighting the importance of clone selection when modifying established panels (**Supplementary Figs. 3B and 4**).

Because unsupervised approaches enable unbiased characterization of phenotypic structure in high-dimensional single-cell datasets, we applied a standardized computational workflow across all acquisitions.^14,15^ After preprocessing and quality control filtering (see Methods), viable CD45⁺CD3⁺ T-cells were exported for UMAP dimensionality reduction and FlowSOM clustering using established high-dimensional cytometry pipelines. To determine whether spectral acquisition improves multidimensional resolution, we compared T-cell landscapes derived from the same samples acquired using CFC and SFC v1, each represented by 20 or 30 FlowSOM-defined clusters (**Fig. 2A–B**). Median-scaled marker expression revealed clear separation of lineage, differentiation, and functional phenotypes, indicating that the panel captures the expected heterogeneity of T-cells under conventional detection, particularly when cells were overclustered (meta = 30) (**Supplementary Fig. 5**). Spectral acquisition maintained the overall topology of the T-cell landscape but generated more sharply defined cluster boundaries and improved separation between clusters across both 20- and 30-cluster FlowSOM configurations, consistent with enhanced resolution obtained through full-spectrum emission modeling (**Fig. 2B**). These improvements were particularly evident along differentiation axes. In CFC data, CCR7, and CD45RA expression delineated canonical naïve (Tn), central memory (Tcm), effector memory (Tem), and terminally differentiated (Temra) subsets in both CD4 and CD8 T-cells (**Fig. 2C and Fig. 2A**). The corresponding SFC v1 projection maintained the same biological organization but exhibited stronger CCR7 and CD45RA gradients and clearer transitional boundaries, indicating improved discrimination of differentiation states (**Fig. 2D**). Notably, CFC failed to clearly separate Tn, Tcm and Tem CD4 T-cells in UMAP space (**Fig. 2A**), a distinction more evident with spectral acquisition (**Fig. 2B**). This reduced ability to clearly separate populations in CFC was reflected by the compressed CCR7 signal distribution compared with SFC (**Fig. 2E**). We next assessed inhibitory receptor resolution, focusing on PD-1, a dim marker particularly sensitive to spreading error.^7,16^ Using FlowSOM-based unsupervised clustering, spectral acquisition increased the proportion of well-resolved PD-1⁺ CD4⁺ T-cell clusters compared with CFC across both 20- and 30-FlowSOM clusters (**Fig. 2F**). Dot plot visualization of PD-1 expression on CD4^+^ T-cells demonstrated a narrower negative baseline and a more distinct positive–negative transition in SFC v1 data, consistent with improved sensitivity for low-intensity markers (**Fig. 2G**).

**Figure 2.**
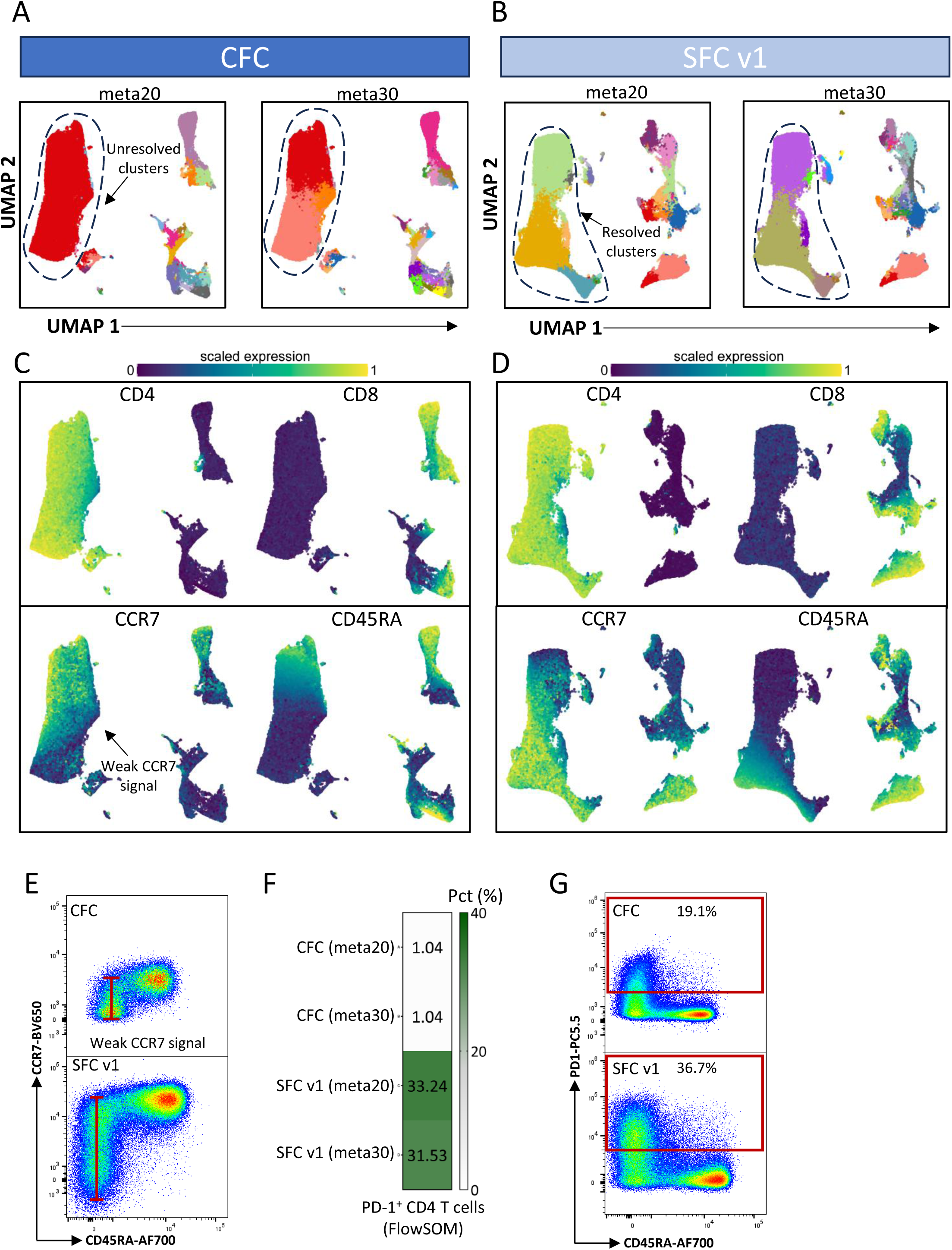
High-dimensional single-cell analysis of total T-cells acquired by CFC or SCFv1. (**A**) Left: UMAP projection showing the distribution of 20 T-cell clusters (meta = 20) derived from CFC acquisition; right: UMAP projection showing the distribution of 30 T-cell clusters (meta = 30) derived from CFC acquisition. (**B**) Left: UMAP projection showing the distribution of 20 T- cell clusters (meta = 20) derived from SFC; right: UMAP projection showing the distribution of 30 T-cell clusters (meta = 30) derived from SFC. (**C**) UMAP visualization of CFC-acquired T-cells colored according to normalized expression of the indicated markers. (**D**) UMAP visualization of SFCv1-acquired T-cells colored according to normalized expression of the indicated markers. (**E**) Representative dot plots showing CCR7 expression in T-cells from both CFC and SFCv1 datasets. (**F**) Heatmap displaying the relative abundance of PD-1⁺ CD4⁺ T-cell clusters identified through unsupervised analysis in CFC and SFC v1 datasets. (**G**) Representative dot plots illustrating manual gating of PD-1⁺ CD4⁺ T-cells in both CFC and SFCv1 datasets.

We next examined the impact of panel optimization within the spectral workflow by comparing SFC v1 and SFC v2 high-dimensional landscapes (**Fig. 3A–B**). Both panels maintained comparable global organization, indicating preservation of biological structure. However, SFC v2 provided clearer separation of residency- and cytotoxicity-associated clusters (C14 and C22) that appeared merged in SFC v1 (C13), suggesting improved marker resolution rather than biological divergence (**Fig. 3A–B**). Although overall results were largely consistent between SFC v1 and SFC v2, subtle differences emerged in a rare cytotoxic CD4⁺ T-cell population (PD-1⁺GZMB⁺)¹⁷ representing approximately 0.72% of total T-cells. Manual gating showed similar frequencies between panels (0.76% vs 0.74%), confirming that the optimized panel did not underestimate this population (**Fig. 3C**). However, only the optimized panel accurately recapitulated this frequency in the unsupervised analysis, indicating improved sensitivity for detecting low-intensity markers (**Fig. 3D**).

**Figure 3.**
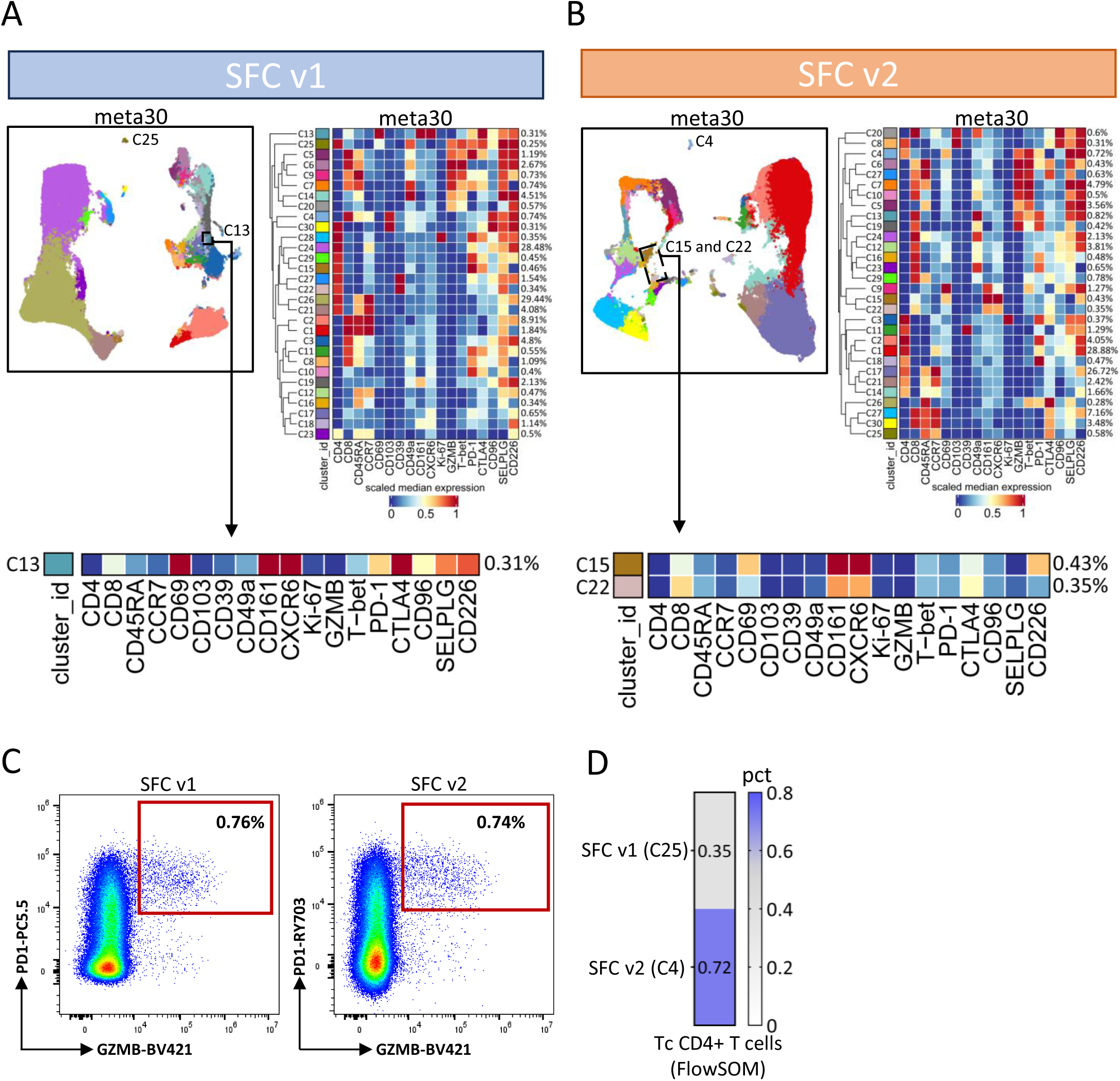
High-dimensional single-cell analysis of total T-cells acquired by SFCv1 or SFCv2. (**A**) Left, UMAP projection illustrating the distribution of T-cells; right, heatmap showing the median scaled expression of selected markers across discrete FlowSOM clusters derived from the SFCv1 dataset. FlowSOM clusters are displayed as rows and markers as columns. The red arrow indicates the presence of cells from the C10 cluster within the C7 cluster island, highlighting the contribution of spillover spreading error to variability in multidimensional data visualization driven by CD49d spread. The boxed area highlights C8 cluster and its phenotypic profile. (**B**) Left, UMAP projection illustrating the distribution of T-cells; right, heatmap showing the median scaled expression of selected markers across discrete FlowSOM clusters derived from the SFCv2 dataset. FlowSOM clusters are displayed as rows and markers as columns. The boxed area highlights C14 and C15 clusters and their phenotypic profile. (**C**) Representative dot plots illustrating manual gating of PD-1⁺ GZMB⁺ CD4+ T-cell in both SFCv1 and SFCv2 datasets. (**D**) Heatmap showing the proportion of PD-1⁺ GZMB⁺ CD4^+^ T-cell clusters identified through unsupervised analysis in SFC v1 and SFCv2 datasets.

Overall, spectral acquisition improved signal-to-noise ratios, staining index values, and multidimensional cluster definition relative to conventional detection while preserving biological organization. Targeted panel optimization further enhanced resolution of co-expressed markers, intracellular transcription factors, and dim checkpoint molecules. However, clone selection may influence quantitative outcomes, underscoring the need for careful reagent validation during panel modification. Collectively, these findings support the combined use of spectral detection and rational panel optimization to improve robustness and sensitivity in high-parameter T-cell phenotyping without fundamentally altering underlying biological interpretation.

## Conclusion

Using a controlled within-instrument comparison on a shared CytoFLEX LX platform, we show that both conventional and spectral flow cytometry reliably capture the overall structure and major populations of the T-cell compartment. CFC remained technically robust and provided consistent identification of principal T-cell subsets, confirming that conventional workflows continue to be suitable for many routine immunophenotyping applications.

However, spectral acquisition provided clear technical advantages, including improved signal-to-noise ratios, higher staining index values, and enhanced resolution of low-intensity and co-expressed markers. These gains translated into sharper multidimensional cluster boundaries, improved discrimination of differentiation states, and more reliable resolution of dim checkpoint molecules such as PD-1. The integration of next-generation fluorophores and targeted panel optimization further reduced spreading effects and enhanced marker separation, particularly for residency-associated and intracellular markers.

Importantly, panel optimization improved unsupervised identification of rare populations without altering their underlying biological frequencies, indicating increased analytical precision rather than biological divergence. At the same time, observed differences related to PD-1 highlighted the strong influence of antibody clone selection on quantitative outcomes, emphasizing the need for careful validation when updating panels.

Overall, while CFC remains effective and appropriate in many experimental contexts, spectral cytometry, especially when combined with new fluorophores and rational panel optimization, provides more precise, higher-quality measurements and improved multidimensional resolution. These improvements enhance the robustness of high-parameter T-cell phenotyping and support the use of optimized spectral workflows in studies requiring sensitive detection of subtle phenotypic differences and rare immune subsets.

## Methods

### Blood collection and isolation of mononuclear cells

All procedures were conducted in accordance with the Declaration of Helsinki and approved by the local institutional ethics committee. Written informed consent was obtained from all donors prior to sample collection. PBMCs were processed and stored following institutional guidelines for the use of human biological materials, and anonymized samples were used for the standardization and optimization of the assays described here. Up to 20 mL of blood was collected from each patient in vacuettes containing ethylenediaminetetraacetic acid (EDTA). Blood was immediately processed. Isolation of PBMCs was performed using Ficoll-Hypaque according to standard procedures.^18^ PBMCs were stored in liquid nitrogen in fetal bovine serum (FBS) supplemented with 10% dimethyl sulfoxide (DMSO). Plasma was stored at −80°C until use.

### Instrument setup and optimization

The CytoFLEX LX and CytoFLEX LX flow cytometers equipped with a mosaic 88 Spectral Detection Module (Beckman Coulter Life Sciences) employ avalanche photodiode detectors (APDs) for fluorescence detection. Instrument setup and quality control were performed using the manufacturer-recommended daily QC workflow with CytoFLEX Daily QC (cat. C65719) and IR (cat. C06147) Fluorospheres verify laser performance, detector sensitivity, and fluidics stability prior to data acquisition. Daily QC and IR ensured consistent target median fluorescence intensity (MFI) values across detectors and confirmed instrument readiness. Detector sensitivity on the CytoFLEX LX instrument is regulated through APD gain settings, which are controlled in instrument-specific units. Default APD gain settings were established by the manufacturer to balance detector sensitivity—defined as the ability to resolve dim signals from background noise—and dynamic range, defined as the ability to accurately capture bright signals without saturation. These settings were optimized based on evaluations of electronic and optical noise, as well as staining index and MFI measurements obtained from PBMCs stained with a broad panel of fluorochrome-conjugated antibodies spanning all excitation lasers and emission ranges (manufacturer data). All data presented in this manuscript were acquired using the default assay gain settings (obtained after QC) for both conventional flow cytometry (CFC) and spectral flow cytometry (SFC) modes. For all samples, populations of interest were confirmed to be fully on scale across all relevant detectors, indicating appropriate gain selection and the absence of detector saturation. For spectral acquisitions performed with the mosaic 88 module, spectral unmixing was conducted using the Beckman Coulter-proprietary Poisson Hybrid algorithm, and detector gains were not modified following daily QC to ensure consistency across experiments. Filter band, laser, and detector information are reported in **Supplementary Fig. 6.**

### Staining protocol

All antibody staining mixes were prepared on the day of use at a final volume of 100 µL per sample. The complete panel was divided into four staining mixes to preserve the optimal fluorochrome concentrations determined by titration (**Supplementary Figs. 7–8**) and to accommodate staining steps requiring sequential incubation. For ex vivo immunophenotyping, cryopreserved blood and tumor samples were thawed in RPMI 1640 with GlutaMAX (Gibco, cat. 61870-010) supplemented with 10% fetal bovine serum (FBS; Euroclone, cat. ECS500L), 1% sodium pyruvate (Gibco, cat. 11360-039), 1% nonessential amino acids (NEAA; Gibco, cat. 11140-035), penicillin–streptomycin (Gibco, cat. 15140-122), 0.1 M 4-(2-hydroxyethyl)-1-piperazineethanesulfonic acid (HEPES; Gibco, cat. 15630-056), and 55 mM β-mercaptoethanol (Gibco, cat. 21985-023), hereafter referred to as R10 medium, and further supplemented with 20 µg/mL DNase I from bovine pancreas (Sigma-Aldrich, cat. DN25G-1G). Cells were washed with PBS and immediately stained with the viability dye Live/Dead NIR876 (Thermo Fisher Scientific, cat. L34980) for 20 min at room temperature in PBS.

After washing with FACS buffer (PBS supplemented with 2% FBS), cells were stained for chemokine receptors (CXCR6 and CCR7) for 30 min at 37°C in a 5% CO_2_ atmosphere. Cells were then washed and stained with a panel of surface monoclonal antibodies (anti-CD45, CD3, CD4, CD8, CD45RA, CD39, CD49a, CD69, CD96, CD103, CD161, CD162, CD226, PD-1, and CTLA-4) for 20 min at room temperature in BV buffer consisting of 50% FACS buffer and 50% Brilliant Stain Buffer (BD, cat. 563794). Intracellular staining for Ki-67, T-bet, and granzyme B (GZMB) was performed following fixation and permeabilization using the FoxP3/Transcription Factor Staining Buffer Set (eBioscience, cat. 00-5523), according to the manufacturer’s instructions, with antibody incubation for 30 min at 4°C. The final dilution of each antibody and its assignment to the corresponding staining mix are detailed in **Supplementary Table 1**.

### Staining index and spreading error calculations

Staining index (SI) and spillover spreading error (SE) were calculated to quantitatively assess marker resolution and the impact of signal spread across detectors. SI was calculated for each marker as the difference between the median fluorescence intensity (MFI) of the positive population and the MFI of the negative population, divided by twice the robust standard deviation of the negative population, using the formula:

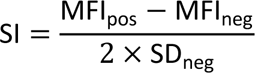

where MFIs and standard deviations were derived from manually gated populations, as indicated. All values used to generate the heatmap shown in Figure 1F are reported in **Supplementary Table 2**.

Spillover spreading error (SE) was calculated following the method described by Nguyen et al.^7^ Briefly, for a given primary fluorochrome and a secondary detector, SE was defined as the difference in the robust standard deviation of the secondary detector signal between the negative population in samples containing the primary fluorochrome and the corresponding negative samples lacking that fluorochrome.^7^ All SI and SE calculations were performed on compensated (CFC) or unmixed (SFC) data using FlowJo-derived statistics, as indicated. Robust statistics were used throughout to minimize the influence of outliers.

### Computational analysis of flow cytometry data

Reference controls were stained in parallel with the corresponding marker-specific staining. Beads were used as reference controls, as this approach reduced autofluorescence-related complexity and improved resolution for both compensation and spectral unmixing. Spectral unmixing was performed using the Beckman Coulter-proprietary Poisson hybrid algorithm, which accounts for the Poisson-distributed nature of signal and noise to manage spillover spreading error (SE) in unmixed data, thereby improving accuracy, particularly for low-intensity signals, in real time. Prior to unsupervised analysis, NxN plots displaying all pairwise combinations of 18 parameters (excluding Live/Dead NIR867, CD3, and CD45) were used to assess the quality of compensation or unmixing (**Supplementary Figs. 9–11**). Compensated or unmixed Flow Cytometry Standard (FCS) 3.0 files were imported into FlowJo (v10.10.0) and subjected to standard gating to remove flow instabilities, doublets, aggregates, and dead cells (**Supplementary Fig. 12**). For each sample acquired by CFC or SFC, 20,000 live CD45⁺CD3⁺ T-cells were selected and imported into R using the flowCore package (v2.10.0). Downstream analyses were performed using the Cytometry dATa anALYSis Tools (CATALYST v1.22.0). This version of CATALYST is based on Bioconductor’s SingleCellExperiment class, which is the standard for high-dimensional single-cell data analysis.^19^ All flow cytometry data were transformed in R using a hyperbolic arcsine transformation [arcsinh(x/cofactor)] with manually defined cofactors, where x represents the measured fluorescence intensity. Cofactors and markers used for unsupervised analyses of CFC, SFCv1, and SFCv2 datasets are reported in **Supplementary Table 3**. Marker distribution quality control (QC) following data pre-processing is shown in **Supplementary Fig. 13**. Clustering and dimensionality reduction were performed using FlowSOM (v2.6.0) and UMAP (uwot v0.2.2), respectively. To facilitate visualization and enable comparison across analyses, the results were displayed using a fixed number of 20 metaclusters for all datasets.

## Data availability

Flow cytometry data from this study are available upon reasonable request; all related inquiries should be directed to the corresponding author.

## Author Contributions

Domenico Lo-Tartaro (D.L.T.): Conceptualization, formal analysis, Investigation, Methodology, Data curation, Validation, Writing – original draft, Writing – review & editing. Kelly Lundsten (K.L.): Support for panel optimization. Anisha Jose (A.J): Writing – Contribution to original draft, Writing – review & editing. Andrea Cossarizza (A.C.) Methodology, Writing – review & editing.

## Acknowledgments

We extend our sincere gratitude to Leonardo Beretta and Fanuel Messaggio (Beckman Coulter Life Sciences) for their valuable technical guidance.

## Supplementary figures

**Supplementary Fig. 1.**
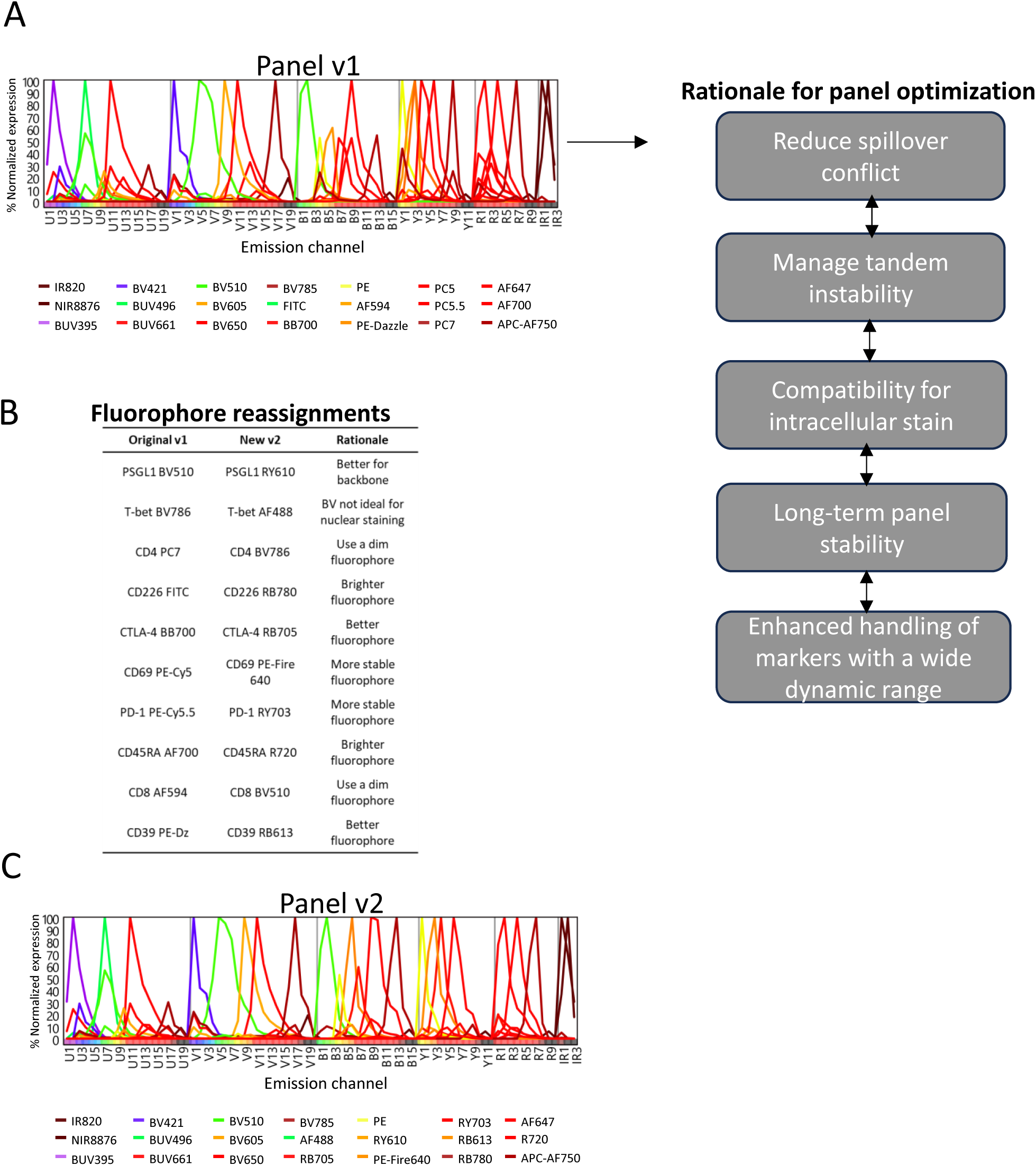
Study overview and panel optimization strategy. (**A**) Left, fluorochrome spectral signatures (normalized emission spectra) of the v1 panel displayed as line plots grouped by emission channel (U, ultraviolet; V, violet; B, blue; Y, yellow; IR, infrared). Right, schematic illustrating the rationale and steps applied for panel optimization.(**B**) Table summarizing fluorochrome reassignments from the original (v1) to the optimized (v2) panel for the indicated antibodies to improve overall data quality.(**C**) Fluorochrome spectral signatures (normalized emission spectra) of the optimized v2 panel shown as line

**Supplementary Fig. 2.**
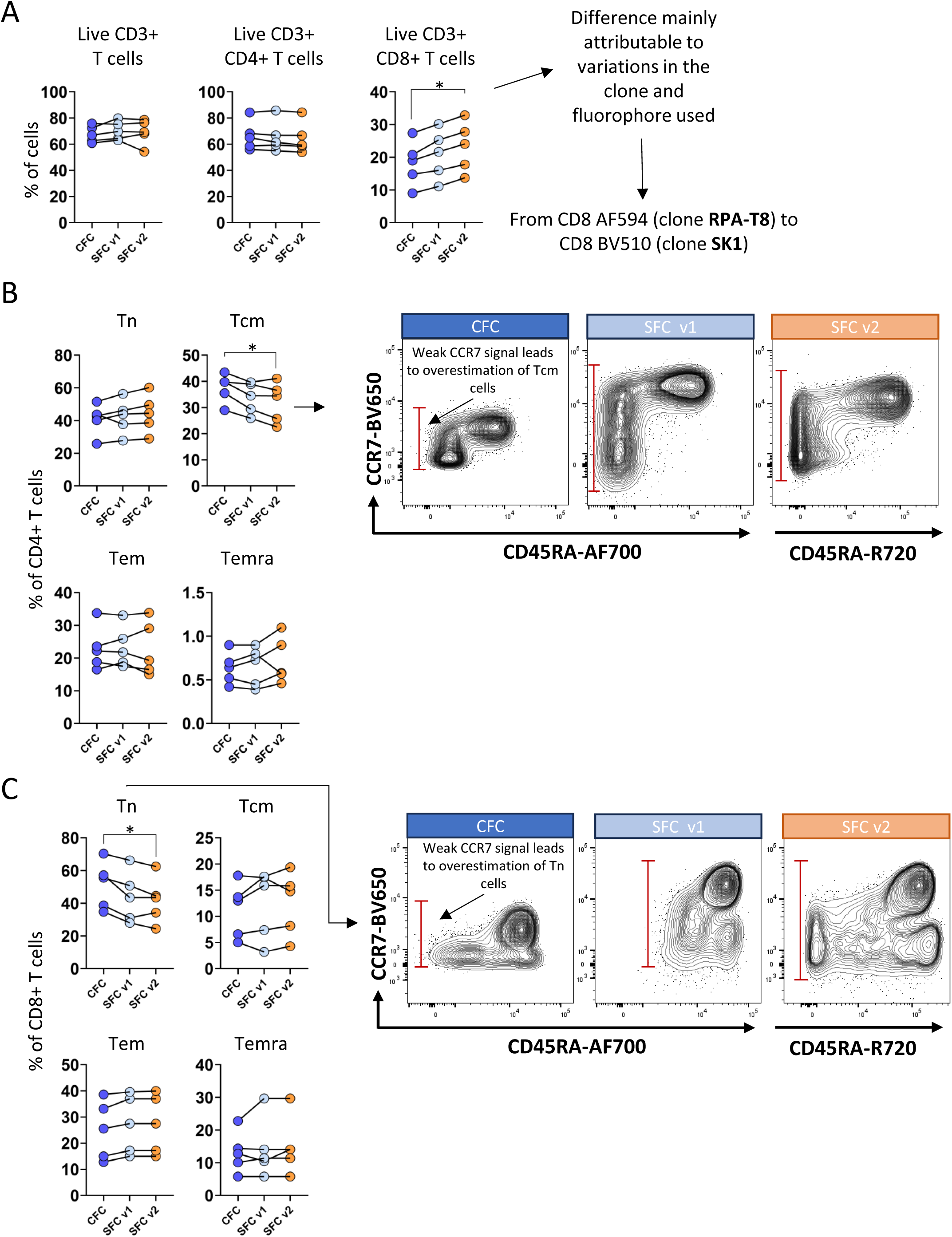
Percentage differences between manually analyzed CFC and SFC datasets for major T cell populations. (A) Scatter plot illustrating percentage changes in CD3⁺, CD4⁺, and CD8⁺ T cells between CFC and SFCv1 or SFCv2.(B) Scatter plot showing percentage changes in naïve (Tn), central memory (Tcm), effector memory (Tem), and effector memory re-expressing CD45RA (Temra) CD4⁺ T cells between CFC and SFCv1 or SFCv2.(C) Scatter plot displaying percentage changes in naïve (Tn), central memory (Tcm), effector memory (Tem), and effector memory re-expressing CD45RA (Temra) CD8⁺ T cells between CFC and SFCv1 or SFCv2.Statistical comparisons were conducted using the Friedman test with Benjamini–Hochberg (BH) correction for false discovery rate (FDR). Statistical significance was defined as FDR < 0.05 (*) and FDR < 0.01 (**).

**Supplementary Fig. 3.**
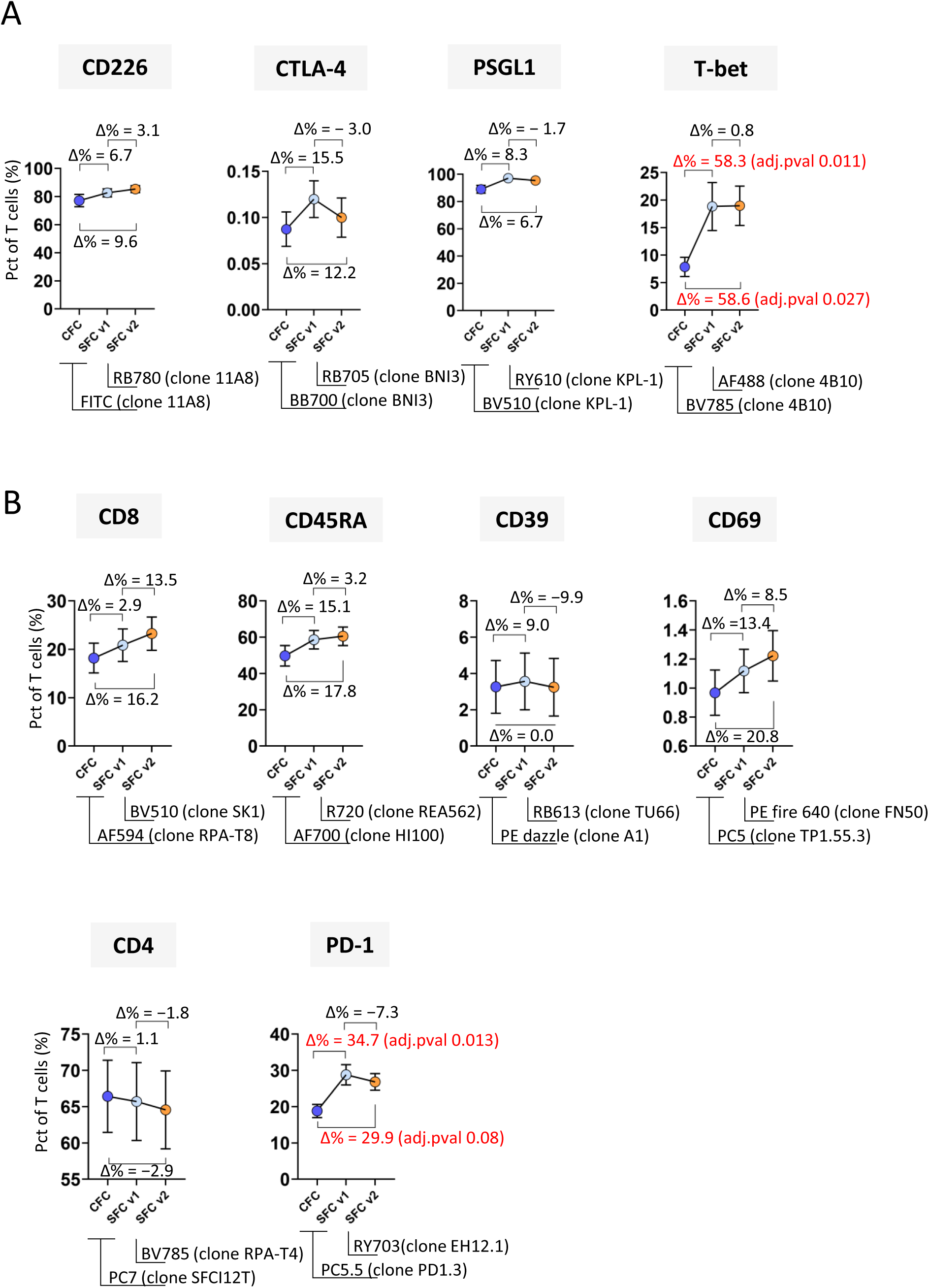
Percentage differences in manually analyzed CFC and SFC data for reassigned antibodies in total of T cells. (**A**) Scatter plot showing the percentage difference (Δ%) of cells positive for the indicated markers within total T cells between CFC and SFCv1 or SFCv2 for antibodies with modified fluorochromes but unchanged clones. CFC-derived percentages were used as the reference. Antibodies showing the largest differences are highlighted in red. (**B**) Scatter plot showing the percentage difference (Δ%) of cells positive for the indicated markers within total T cells between CFC and SFCv1 or SFCv2 for antibodies in which both fluorochrome and clone were modified. Percentages derived from CFC were used as the reference. Antibodies exhibiting the greatest differences are highlighted in red. The reported Δ% values represent the differences between the mean values obtained from the five analyzed samples.

**Supplementary Fig. 4.**
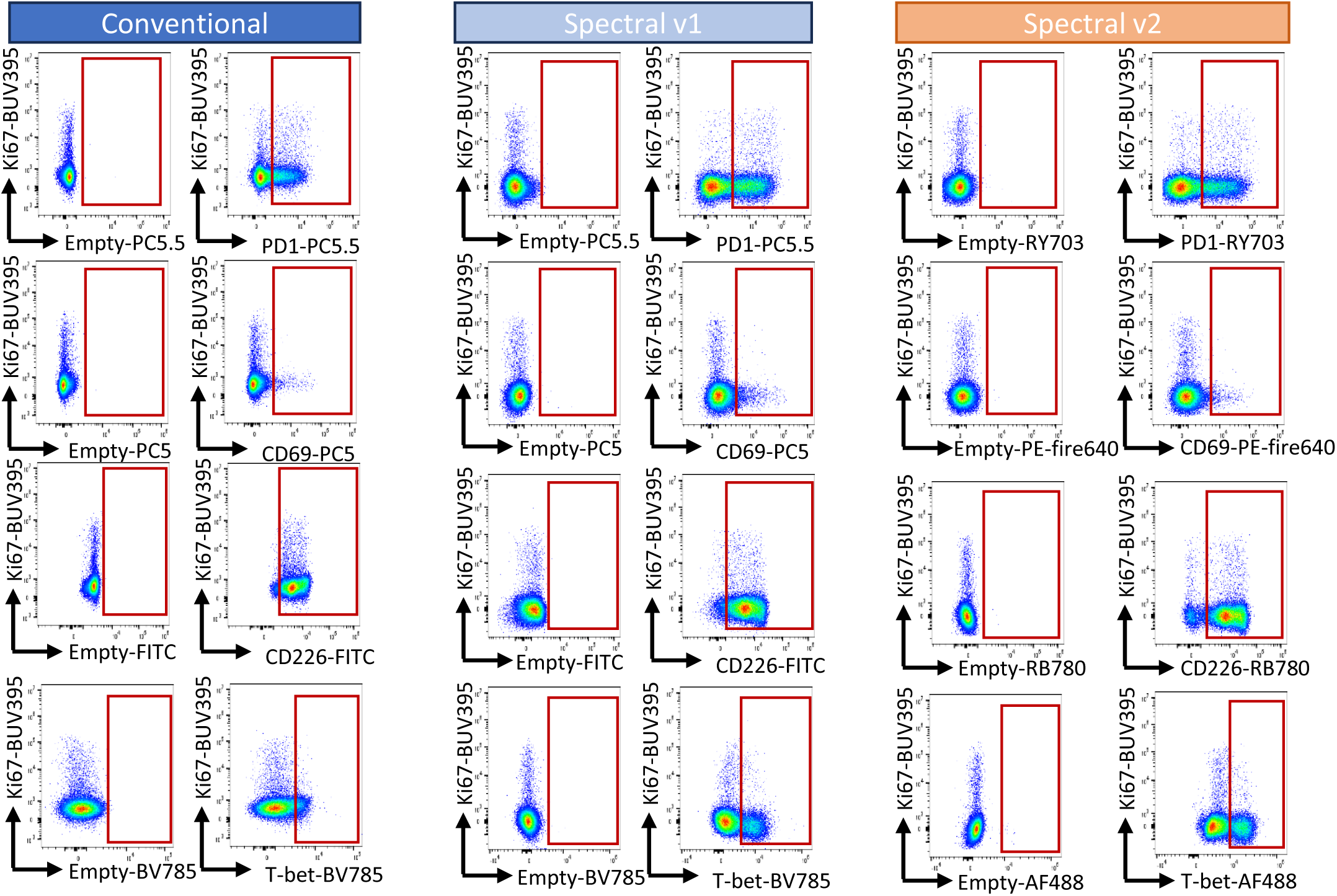
Marker positivity of antibodies exhibiting the largest differences between CFC and SFC analyses. Dot plots display markers with the highest percentage variation (Δ%), indicated in red in **Supplementary** Fig. 2. The empty channel represents fluorescence-minus-one (FMO) controls used to establish positivity thresholds.

**Supplementary Fig. 5.**
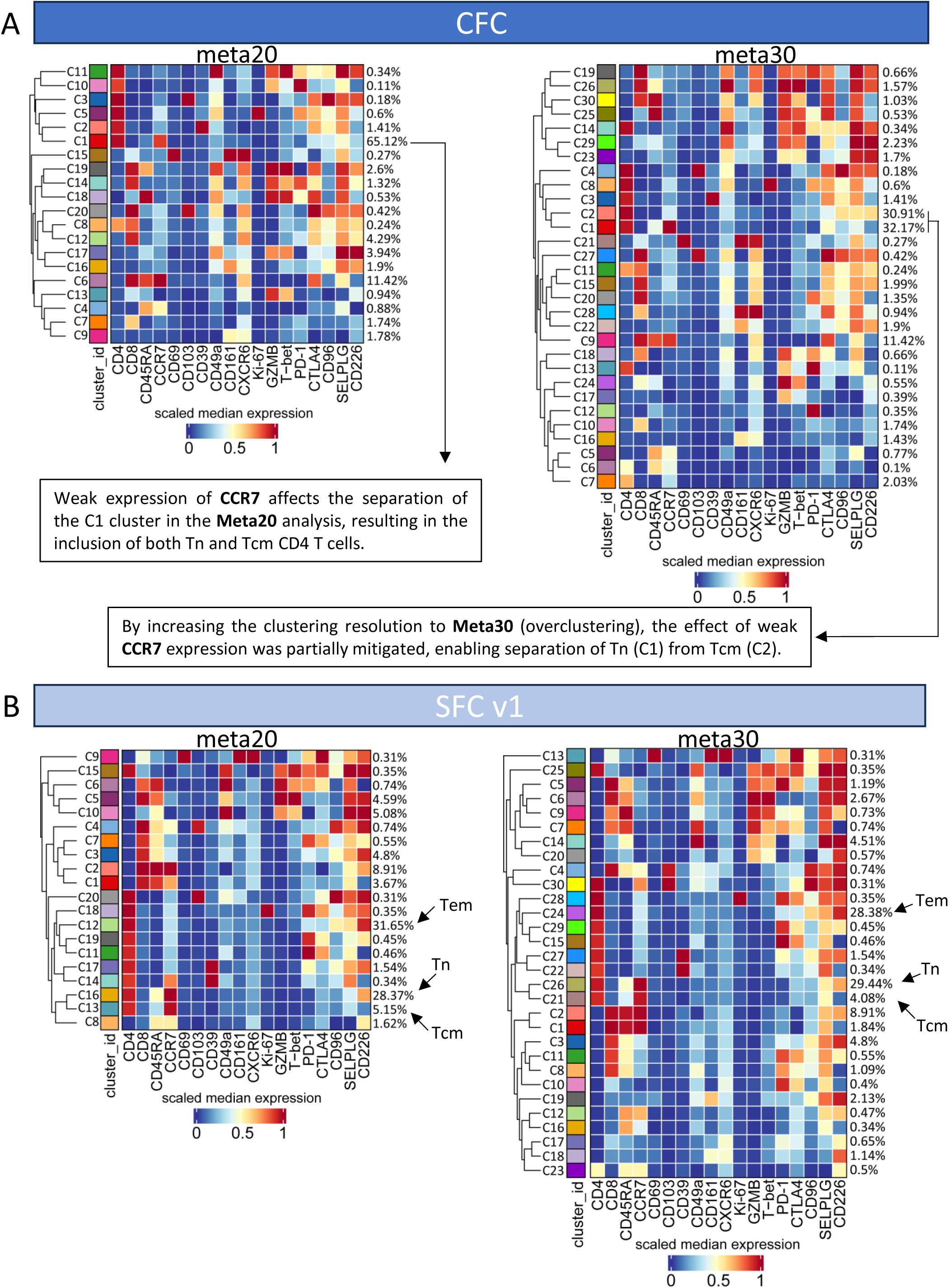
FlowSOM clustering of CFC and SFC v1 samples. (A) Heatmap showing the median scaled expression of selected markers across FlowSOM clusters from the CFC dataset at two metaclustering resolutions (meta20 and meta30). Clusters are shown as rows and markers as columns.(B) Heatmap showing the median scaled expression of selected markers across FlowSOM clusters from the SFC v1 dataset at two metaclustering resolutions (meta20 and meta30). Clusters are shown as rows and markers as columns.

**Supplementary Fig. 6.**
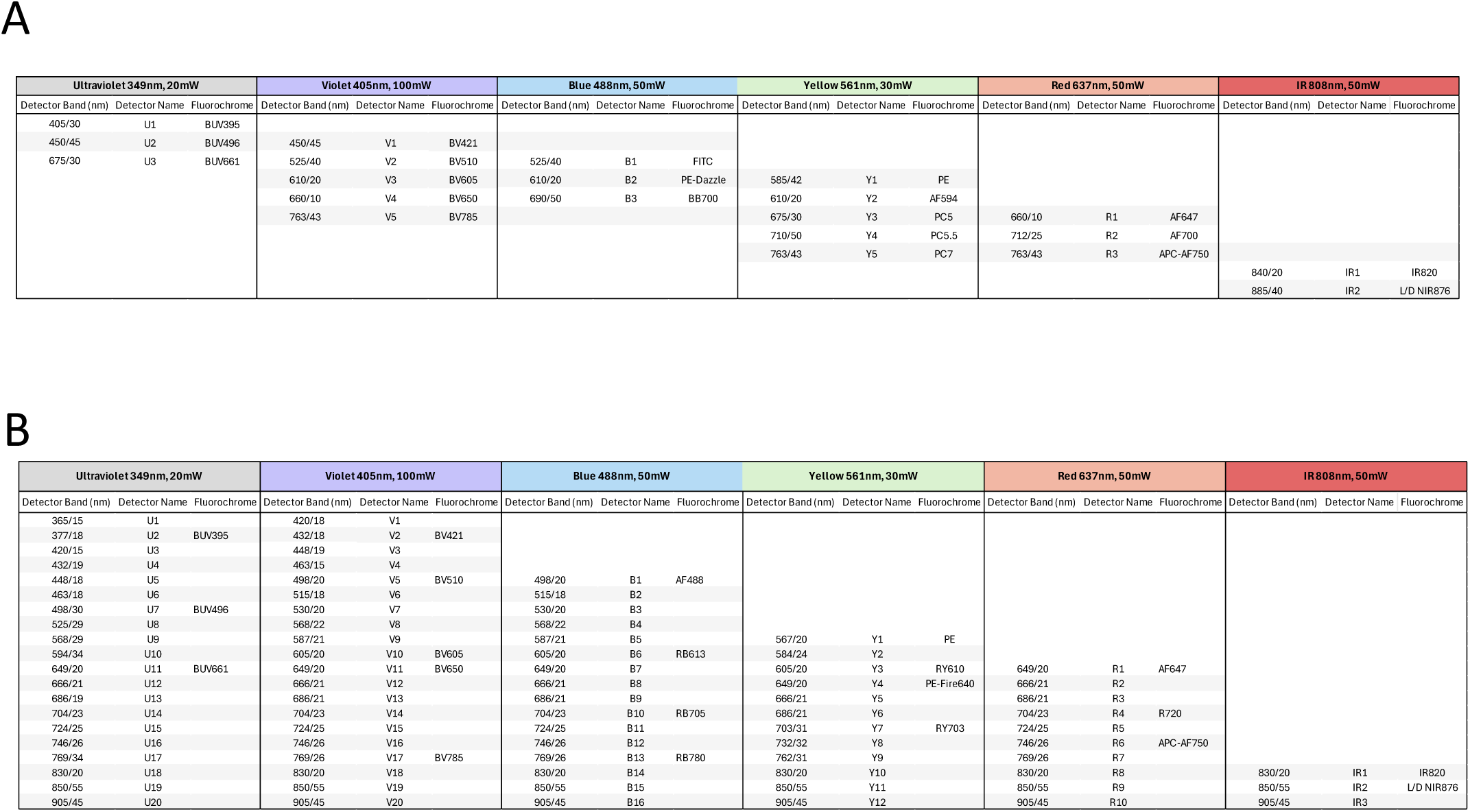
Instrument configuration. Filter band, laser, and detector information are shown for all spectral detectors on the Beckman Coulter CytoFLEX LX and CytoFLEX LX equipped with Mosaic 88, together with the final 21 fluorochromes used in this study and their representative detectors, which for most fluorochromes correspond to the peak detector. The first column reports the effective detection band for each spectral detector, expressed as effective center wavelength (CWL) and effective bandwidth (BW) in nanometers, across all six lasers used for spectral detection. Detector names indicate their relative position within each laser’s detector array and their approximate center wavelength. Effective CWL and BW values are determined by the configuration of bandpass filters in the optical path. Non-spectral detectors, including scatter and imaging channels, are not shown. (**A**) Instrument configuration of the CytoFLEX LX used in conventional flow cytometry mode (CFC).(**B)** Instrument configuration of the CytoFLEX LX equipped with Mosaic 88 used in spectral flow cytometry mode (SFC).

**Supplementary Fig. 7.**
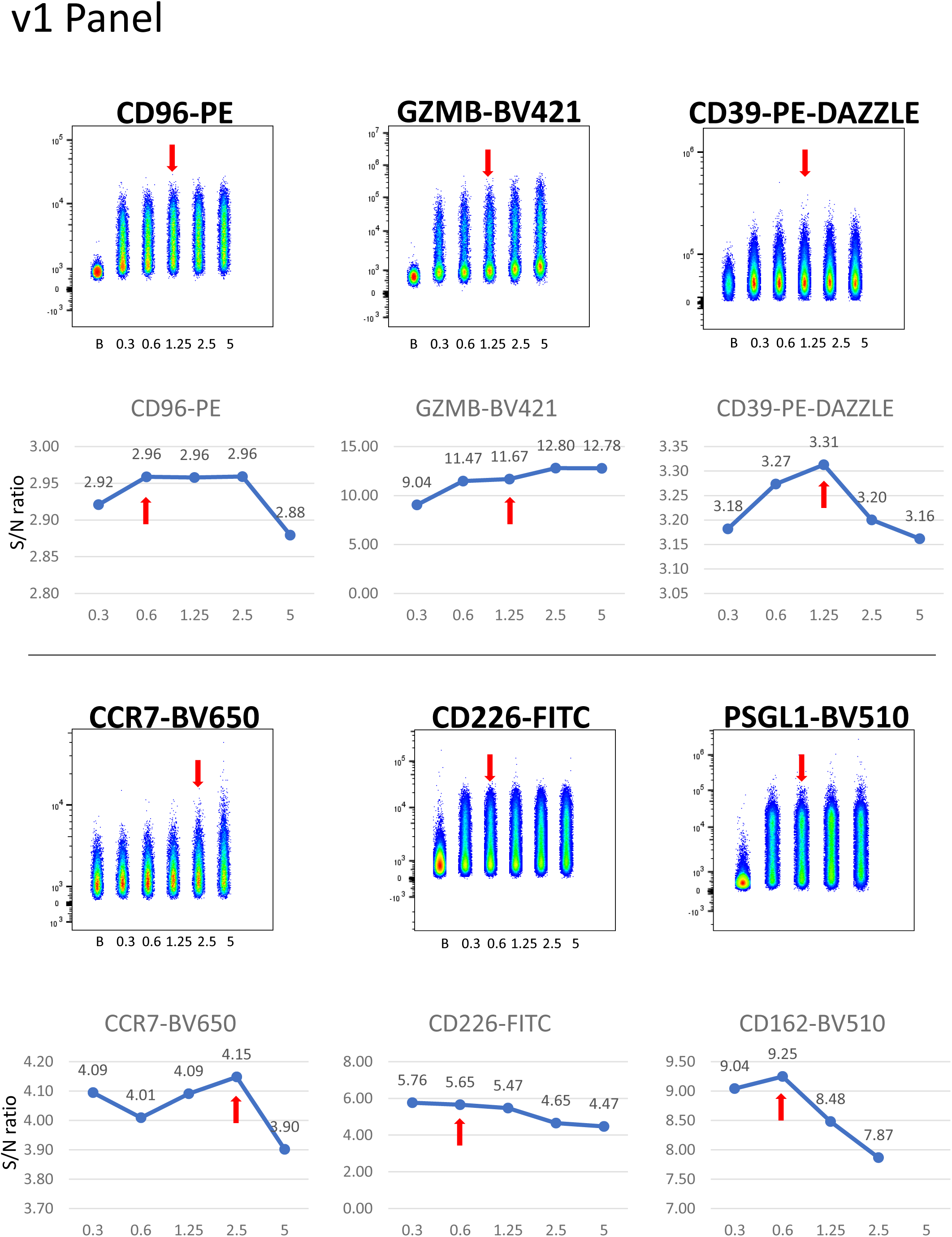
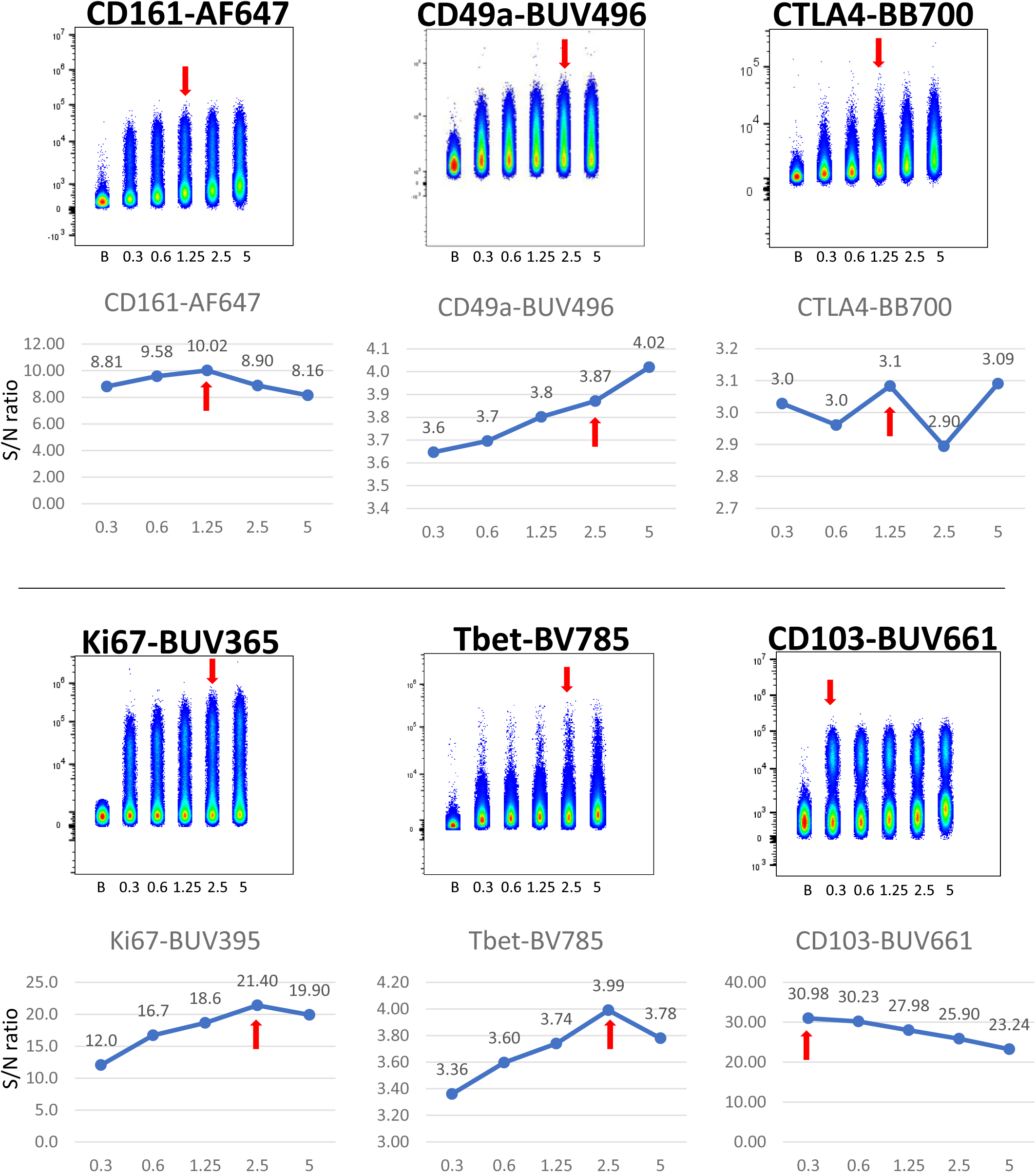
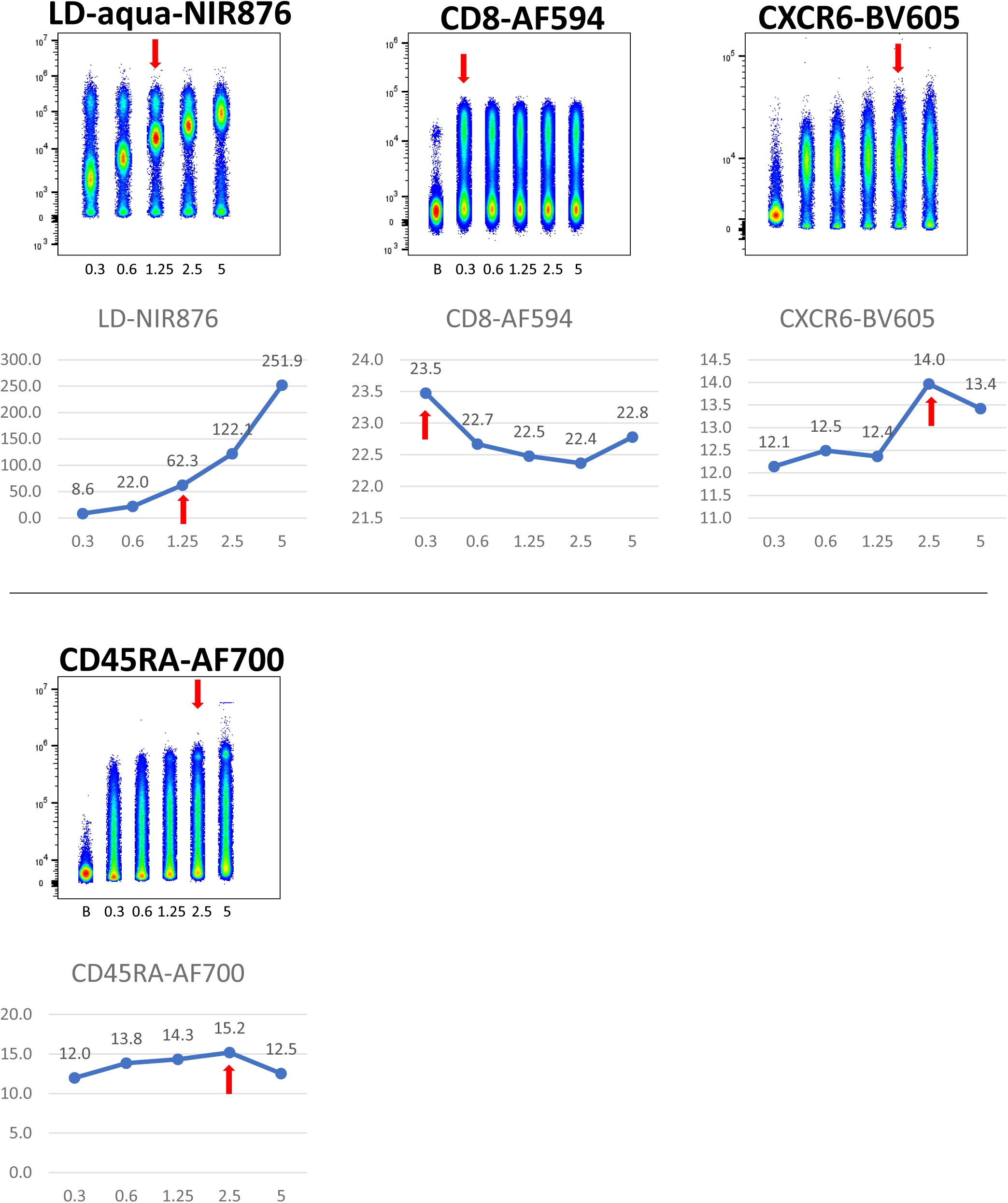
Titrations for fluorochrome-antibody conjugates used in the final 21-color original panel (v1). Titrations were performed on cryopreserved PBMCs using serial dilutions ranging from 5 µL/test to 0.3 µL/test, unless otherwise indicated, and samples were acquired on a CytoFLEX LX + Mosaic88. Antibody titrations were conducted on pre-gated T cells, which were identified by prestaining with CD3-FITC or CD3-PB or CD3-PE—selected to minimize spectral overlap with the antibody under titration—together with Live/Dead NIR876. Individual FCS files were concatenated for visualization and signal to noise ratio (MFI+ / MFI−) was calculated to optimal concentration (red arrow). Antibodies from Beckman Coulter were used at the maximum recommended titer, as reducing the concentration resulted in excessive signal loss (data not shown).

**Supplementary Fig. 8.**
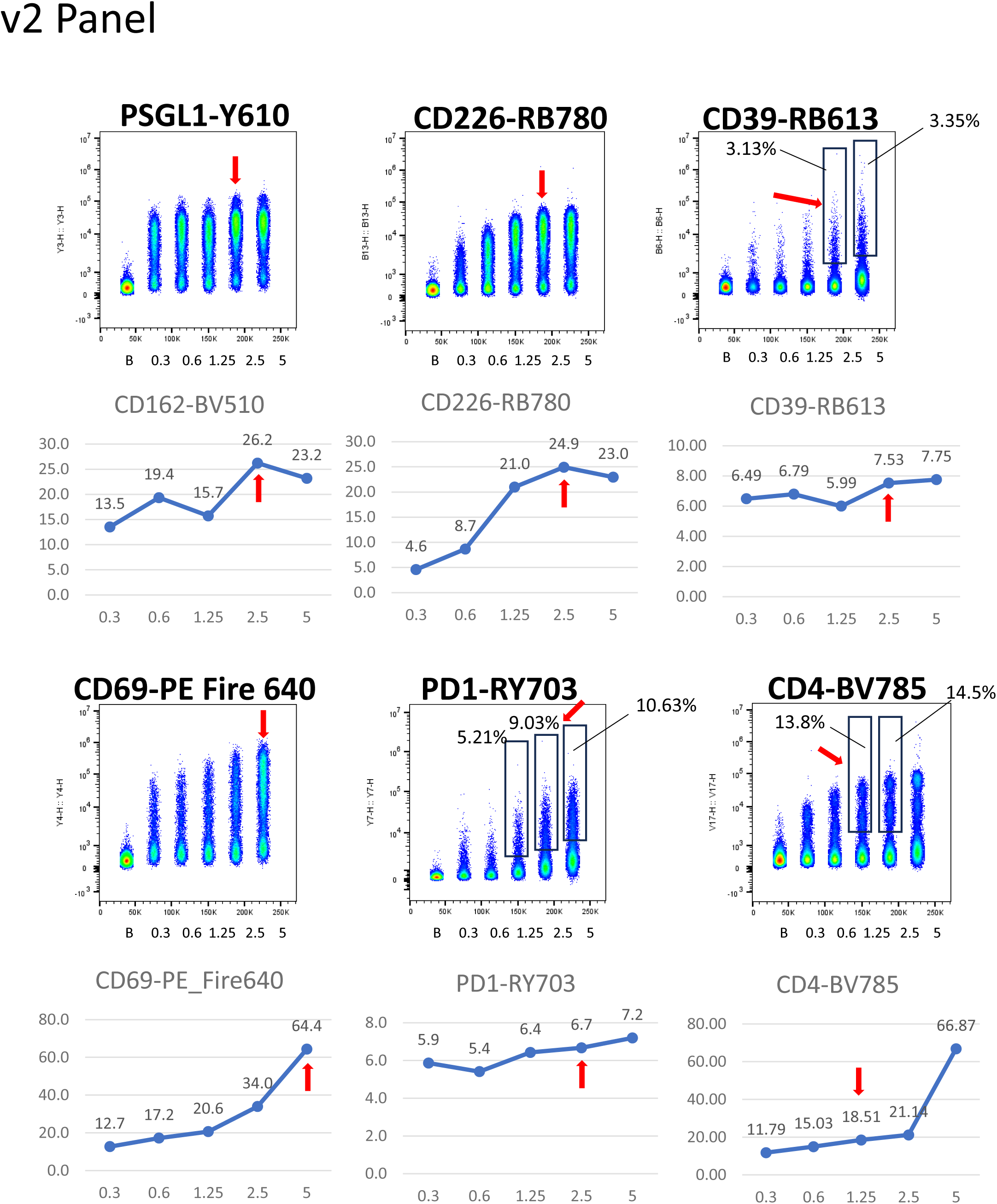
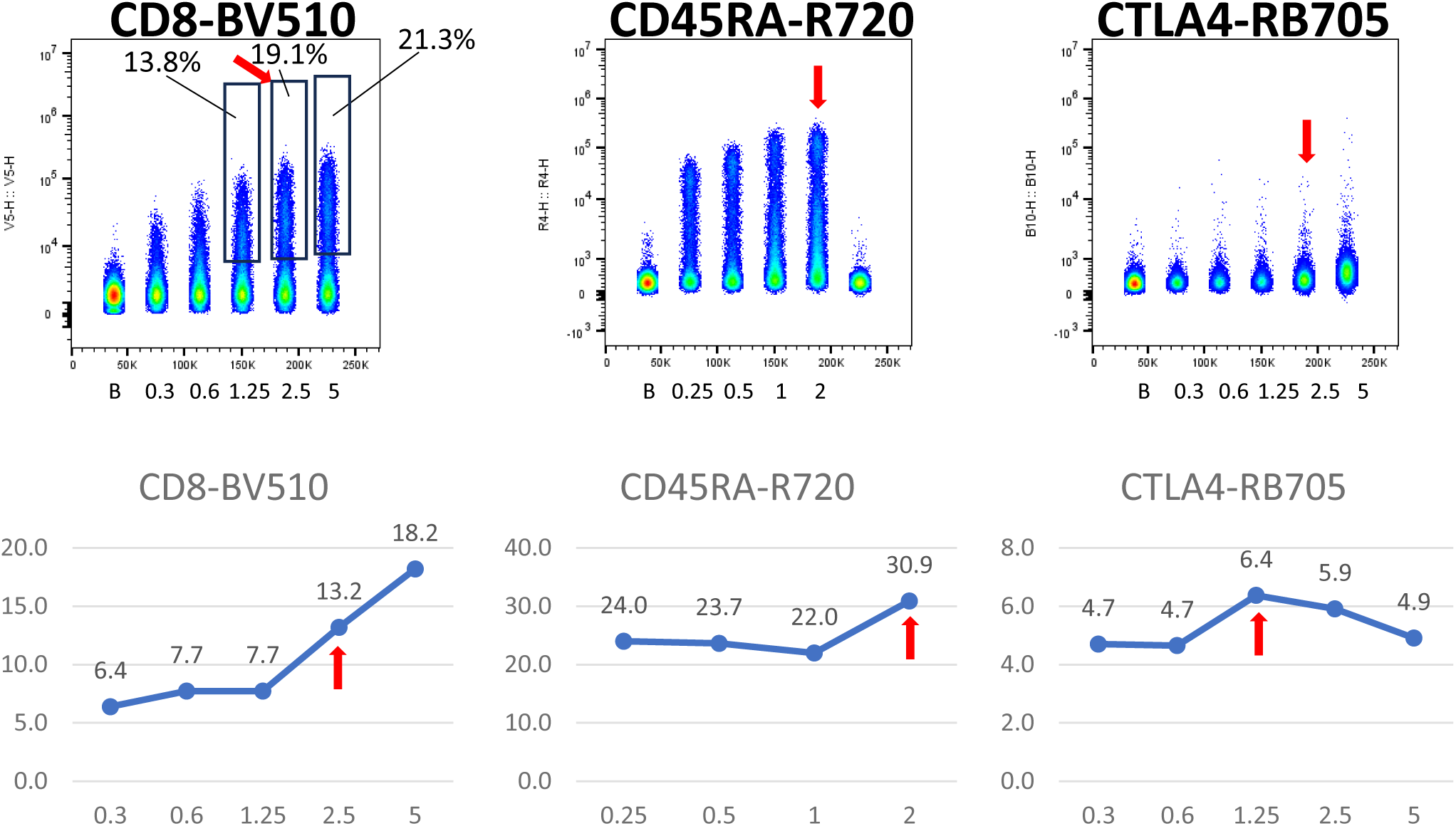
Titration of newly introduced fluorochrome–antibody conjugates selected during panel optimization for the final 21-color panel (v2). Titrations were performed on cryopreserved PBMCs using serial dilutions ranging from 5 µL/test to 0.3 µL/test, unless otherwise indicated, and samples were acquired on a CytoFLEX LX + Mosaic88. Antibody titrations were conducted on pre-gated T cells, which were identified by prestaining with CD3-FITC or CD3-PB or CD3-PE—selected to minimize spectral overlap with the antibody under titration—together with Live/Dead NIR876. Individual FCS files were concatenated for visualization and signal to noise ratio (MFI+ / MFI−) was calculated to optimal concentration (red arrow).

**Supplementary Fig. 9.**
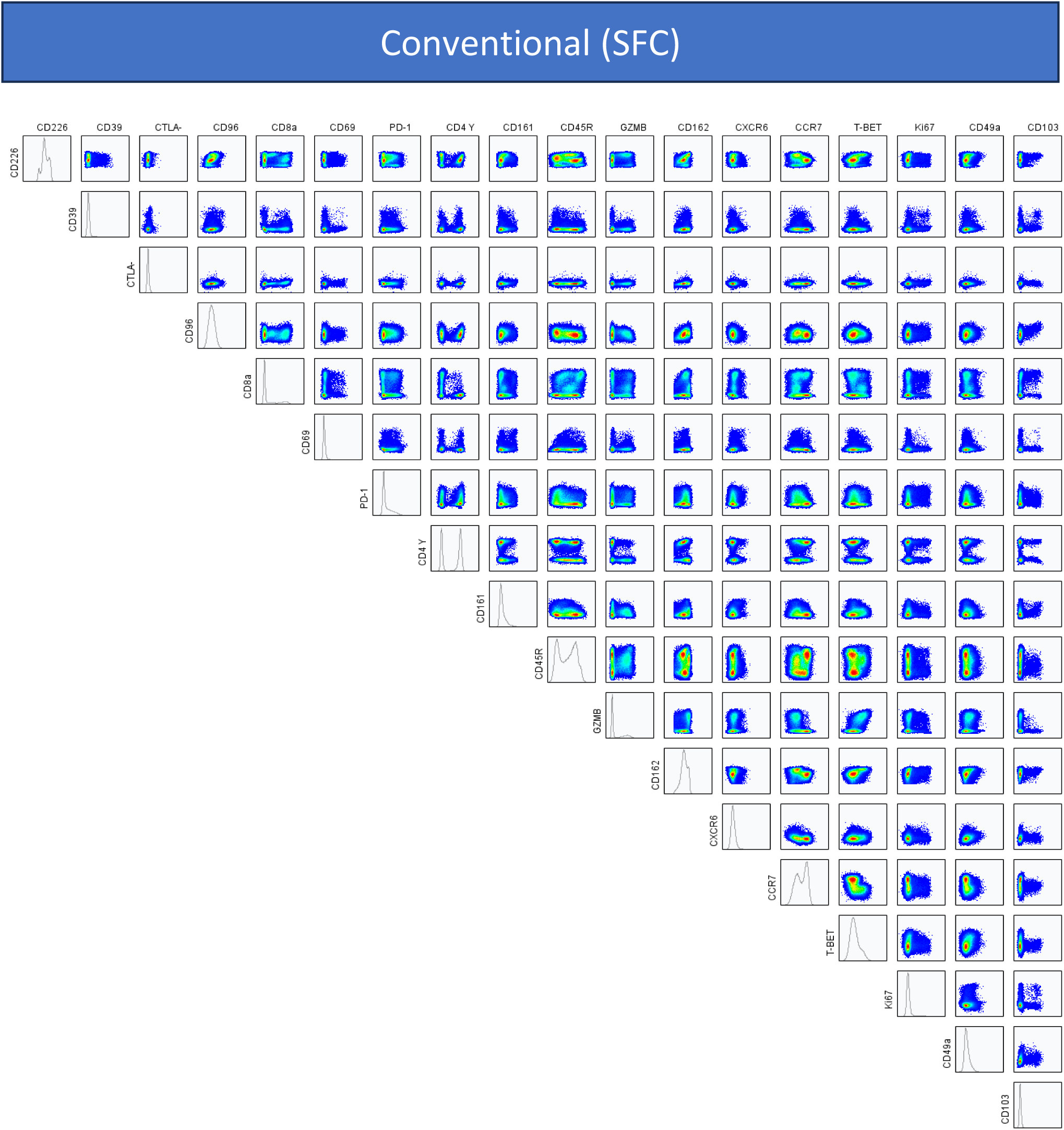
Compensation quality control (QC). NxN plot showing all pairwise combinations of 18 parameters (excluding Live/Dead NIR867, CD3, and CD45) in exported T cells following manual preprocessing (see Supplementary Fig. 12).

**Supplementary Fig. 10.**
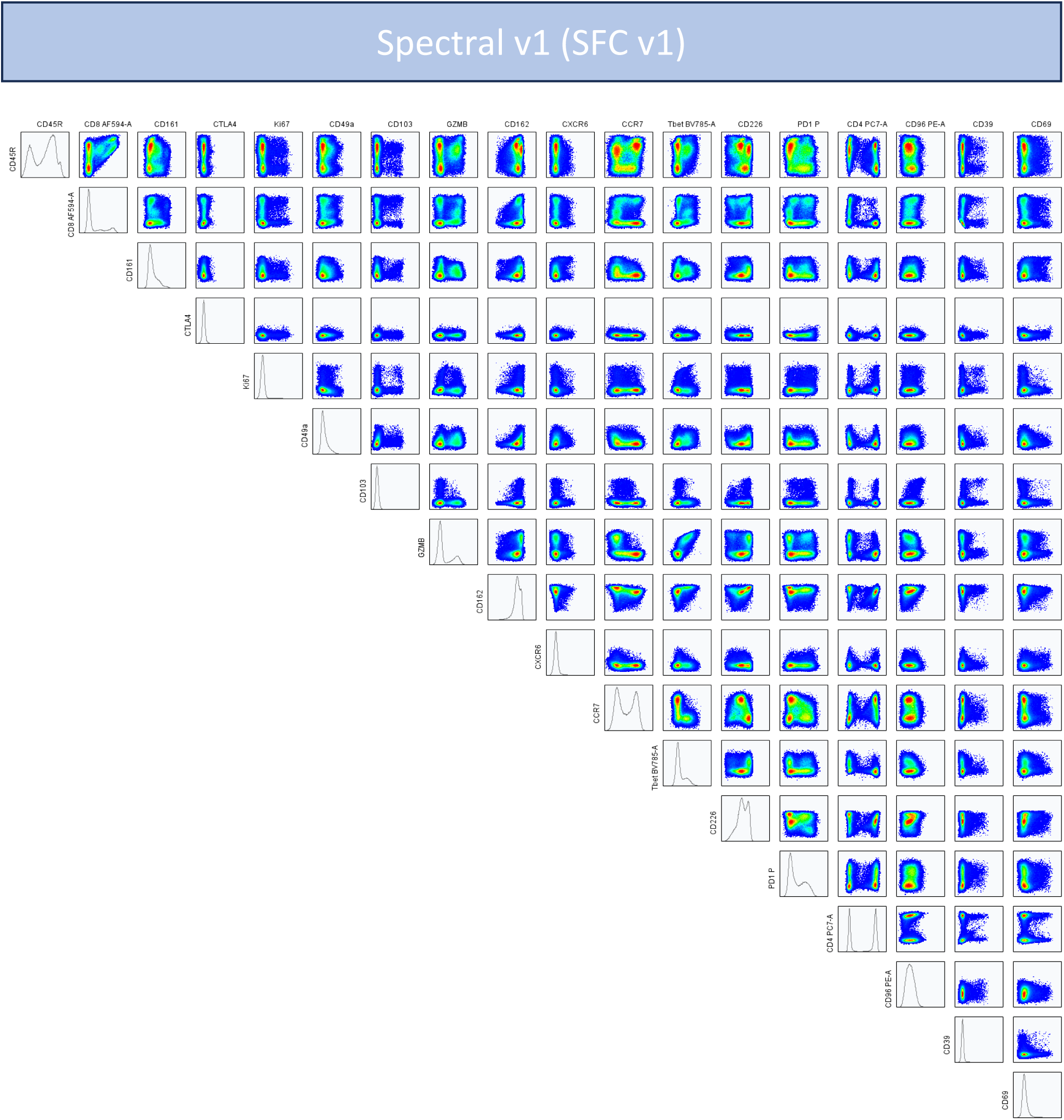
Unmixing quality control (QC) of SFC v1 .fcs file. NxN plot showing all pairwise combinations of 18 parameters (excluding Live/Dead NIR867, CD3, and CD45) in exported T cells following manual preprocessing (see **Supplementary Fig. 12**).

**Supplementary Fig. 11.**
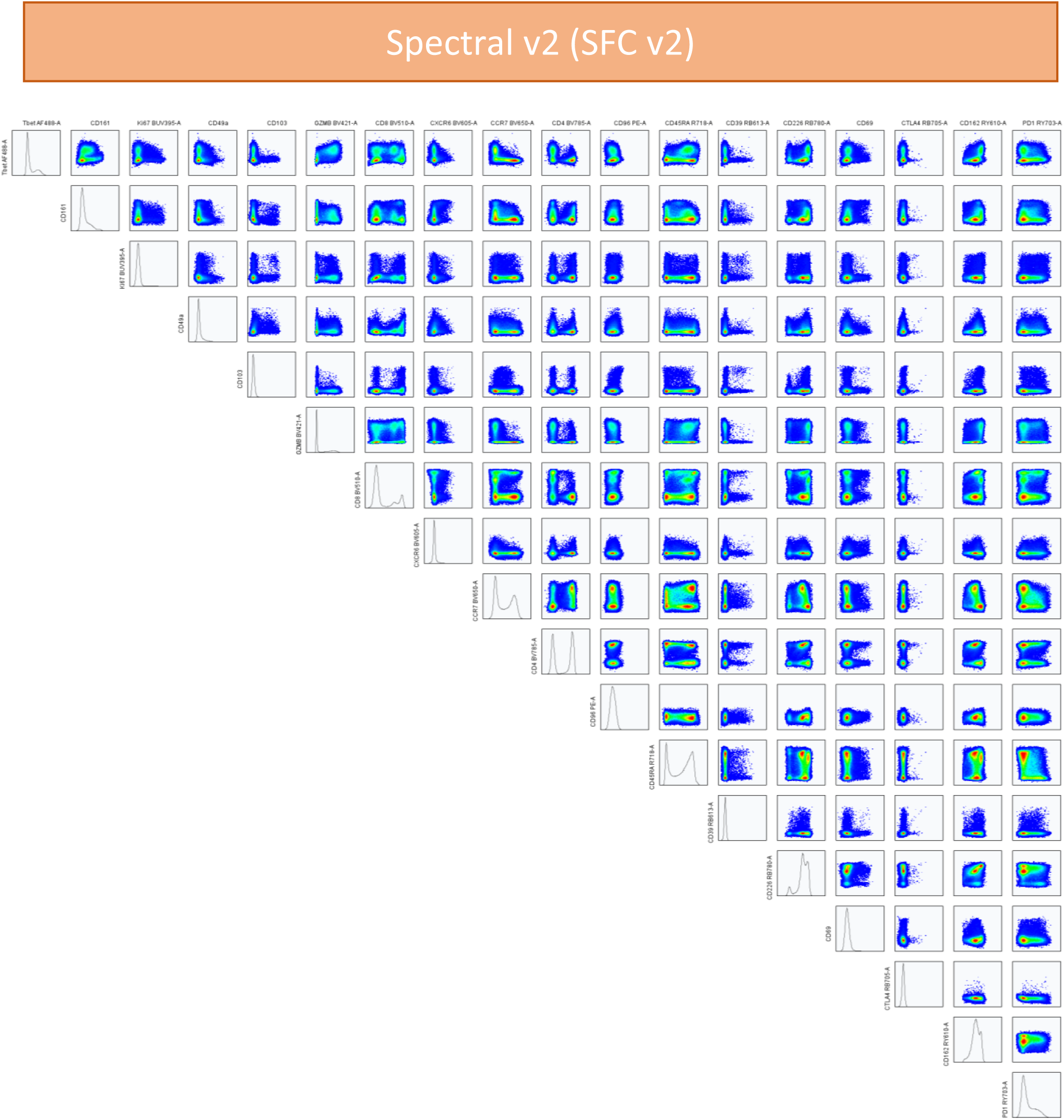
Unmixing quality control (QC) of SFC v2 .fcs file. NxN plot showing all pairwise combinations of 18 parameters (excluding Live/Dead NIR867, CD3, and CD45) in exported T cells following manual preprocessing (see **Supplementary Fig. 12**).

**Supplementary Fig. 12.**
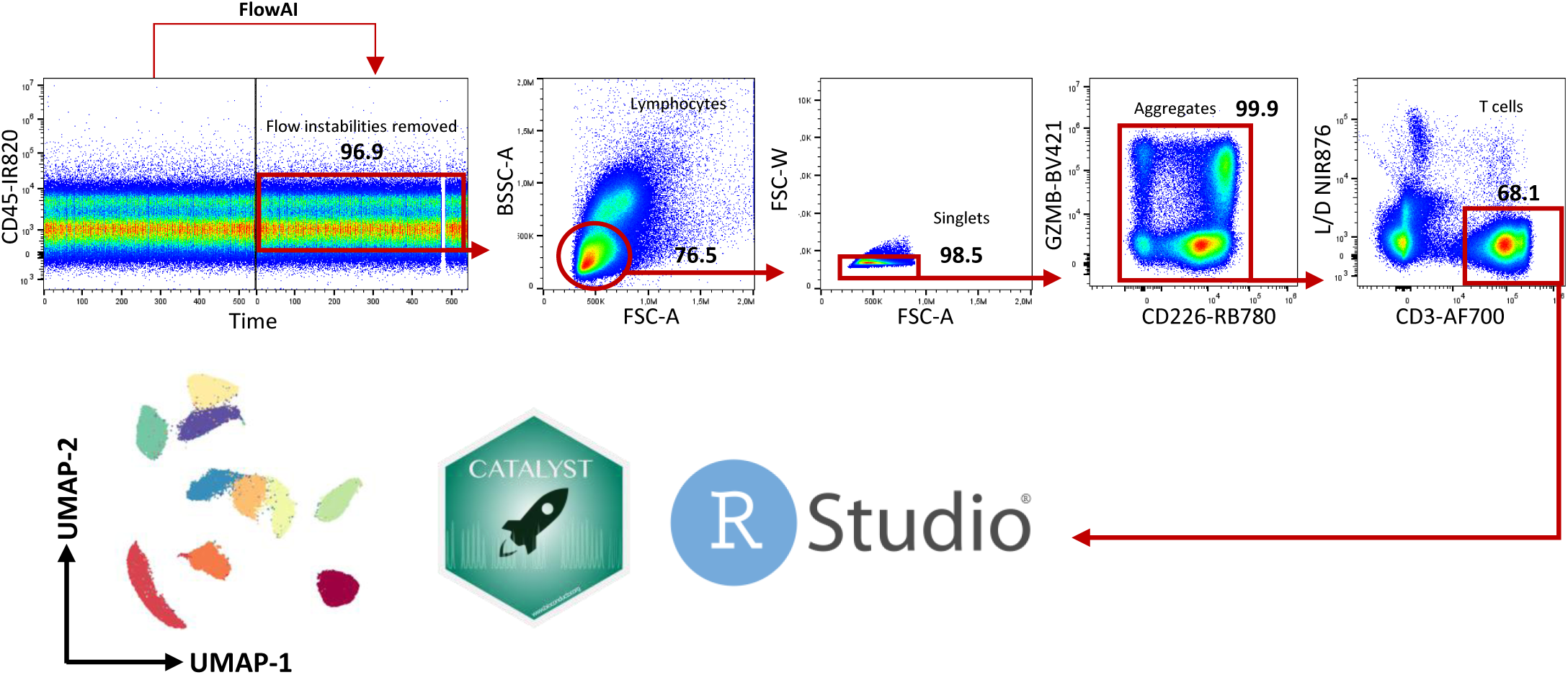
Flow cytometry data preprocessing prior to high-dimensional analysis. Gating strategy illustrating the manual preprocessing workflow used to isolate viable CD45⁺CD3⁺ T cells. All .fcs files were initially imported into FlowJo and processed with FlowAI to remove outliers and unstable events generated during acquisition. Filtered files were then manually inspected to exclude doublets, dead cells, debris, and aggregates. T cells were subsequently selected, exported, and uploaded into R for analysis using the CATALYST package.

**Supplementary Fig. 13.**
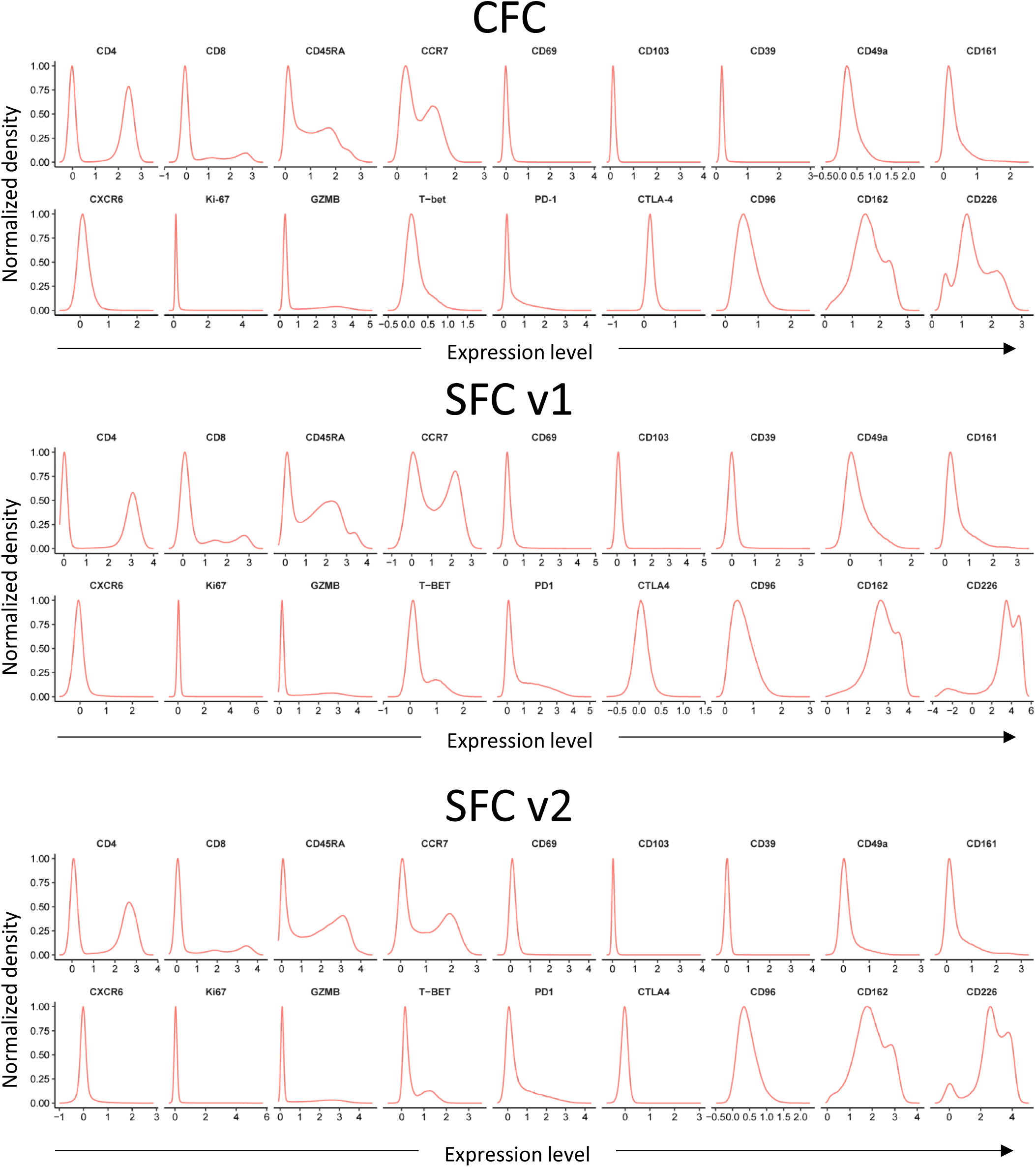
Marker distribution quality control (QC). Histogram plots illustrating marker expression profiles stratified by CFC, SFCv1, and SFCv2. The y-axis represents normalized density, while the x-axis shows marker expression levels expressed as arcsinh-transformed intensity.

